# Ventral pallidum projections to the ventral tegmental area reinforce but do not invigorate reward-seeking behavior

**DOI:** 10.1101/2023.05.22.541796

**Authors:** Dakota Palmer, Christelle A. Cayton, Alexandra Scott, Iris Lin, Bailey Newell, Morgan Weberg, Jocelyn M. Richard

## Abstract

Reward-predictive cues acquire motivating and reinforcing properties that contribute to the escalation and relapse of drug use in addiction. The ventral pallidum (VP) and ventral tegmental area (VTA) are two key nodes in brain reward circuitry implicated in addiction and necessary for the performance of cue-driven behavior. Evidence suggests that VP neurons projecting to the VTA (VP→VTA) promote cue-induced reinstatement of drug-seeking, but the mechanisms by which these neurons do so are undefined. In addition, the role of these neurons in the pursuit of non-drug reward is not known. In the current study, we used *in vivo* fiber photometry and optogenetics to record from and manipulate VP→VTA in rats performing a discriminative stimulus task (DS task) with sucrose reward to determine the fundamental role these neurons play in invigoration and reinforcement by reward and associated discriminative cues. We find that VP→VTA neurons are selectively active during reward consumption, that optogenetic stimulation of these neurons paired with reward consumption biases choice, and that VP→VTA optogenetic stimulation is reinforcing. Critically, we found no significant encoding of cue-elicited reward-seeking vigor and acute optogenetic stimulation of these neurons paired with cue onset did not enhance the probability or vigor of reward-seeking. Our results suggest that VP→VTA neurons are active during the consumption of natural reward and that this activity reinforces seeking behavior.

## INTRODUCTION

Animals use sensory cues associated with threats and rewards to make predictions and inform decision-making. In addition to their predictive utility, reward-predictive cues can acquire incentive motivational value which enables them to capture attention, enhance desire, and powerfully bias animal behavior in favor of reward pursuit and consumption (Robinson and Berridge, 2000). While evolutionarily advantageous in naturalistic settings, incentive motivational processes contribute to maladaptive reward-seeking and consumption characteristic of addiction (Ferguson and Shiffman, 2009; Preston et al., 2018). Characterizing the neurobiological mechanisms by which reward-predictive cues motivate and reinforce seeking behavior is therefore critical.

The ventral pallidum (VP) is a basal forebrain nucleus crucial in the performance of cue-driven behavior (Smith et al., 2009; Root et al., 2015; Soares-Cunha and Heinsbroek, 2023). Recent circuit-specific manipulations have identified functionally divergent roles of VP neurons based on output pathway (Knowland et al., 2017; Faget et al., 2018; Heinsbroek et al., 2020; Prasad et al., 2020; Stephenson-Jones et al., 2020). VP neurons projecting to the ventral tegmental area (VTA) appear specifically to promote cue-driven behavior, including cue-induced reinstatement of cocaine seeking (Mahler and Aston-Jones, 2012; Mahler et al., 2014) and context-induced reinstatement of alcohol seeking (Prasad and McNally, 2016). However, the underlying neurobiological and psychological mechanisms by which these neurons mediate cue-elicited reward seeking are unknown. Prior extracellular recordings identified VP neurons which respond to discriminative cues and predict the vigor of animals’ reward-seeking (Richard et al., 2016, 2018; Lederman et al., 2021). Parallel work has demonstrated that VTA dopaminergic signaling promotes effortful responding to discriminative cues (Yun, 2004; Fischbach-Weiss et al., 2018). In addition to vigor-encoding neurons, VP contains single-units which encode reward-prediction error signals (Tian et al., 2016; Ottenheimer et al., 2020) resembling those canonically associated with VTA dopamine signaling and reinforcement learning (Arsenault et al., 2014; Watabe-Uchida et al., 2017; Lerner et al., 2021). Thus, evidence exists suggesting that VP and VTA support both cue-elicited invigoration and/or reinforcement.

While VP and VTA are both crucial nodes in reward circuitry implicated in addiction, the specific role of VP neurons projecting to the VTA (VP→VTA) is poorly defined. Prior literature probing these neurons during cue-elicited behavior has largely focused on their contribution to cue-induced reinstatement of drug-seeking, in which Pavlovian cues are presented in response to the action previously associated with drug delivery. The translational relevance of such paradigms has been debated in part because those who suffer from addiction often passively encounter cues outside of their control, including discriminative cues signaling reward availability (Troisi, 2013; Namba et al., 2018; Lay and Khoo, 2021). In the current study, we used *in vivo* fiber photometry and optogenetics to record from and manipulate VP→VTA neurons in rats performing a discriminative stimulus task (DS task) (Nicola et al., 2004; Richard et al., 2016, 2018) with sucrose reward to determine the fundamental roles these neurons play in invigoration and reinforcement by reward and associated discriminative cues. For optogenetic manipulations, we compared manipulation of VP→VTA to manipulation of another major output of the VP: VP neurons projecting to the mediodorsal thalamus (VP→mdThal) (Young et al., 1984; Root et al., 2015; Engeln et al., 2022; Soares-Cunha and Heinsbroek, 2023). We find that VP→VTA neurons are selectively active during reward consumption, that optogenetic stimulation of these neurons paired with reward consumption biases choice, and that VP→VTA optogenetic stimulation is reinforcing. Critically, we found no significant encoding of cue-elicited reward-seeking vigor and acute optogenetic stimulation of these neurons paired with cue presentations did not enhance the probability or vigor of reward-seeking. Our results suggest that VP→VTA neurons are active during the consumption of natural reward and that this activity reinforces seeking behavior.

## MATERIALS AND METHODS

### Subjects

Male and female adult Long-Evans rats (Envigo, n=50, 29 female) weighing 250-274g upon arrival were used. Rats were initially pair-housed under a standard 12:12 Light:Dark cycle on ventilated racks. Following surgery, rats were housed singly. Water and food were available *ad-libitum* throughout most of the experiments, except during the choice task experiment. During this experiment, rats were mildly food restricted to 90% of their body weight. All behavioral testing occurred during the light phase. All experimental procedures were approved by the Institutional Animal Care and Use Committee at the University of Minnesota and were carried out in accordance with the guidelines on animal care and use of the National Institutes of Health of the United States.

### Experimental design

Each experiment described incorporated within- and between-subject controls to minimize the influence of confounding variables. For photometry recordings, motion artifacts were removed within-subject using a 405nm isosbestic, calcium-independent signal recorded in parallel to the 465nm calcium-dependent signal (Zalocusky et al., 2016). Photometry signals were z-scored on a trial-by-trial basis to normalize between-subjects. Behavioral tasks included control stimuli and operands in addition to reward-associated stimuli and operands. The DS task included presentations of a neutral control cue (NS) as well as the DS to account for general impacts of auditory stimuli on neural activity and behavior. Cue identity of the DS and NS was counterbalanced between-subjects. To assess specificity of manipulating distinct VP subpopulations during similar epochs, we performed similar optogenetic manipulations of VP neurons projecting to the mediodorsal thalamus (VP→mdThal). DS task sessions with optogenetic manipulations contained laser-paired as well as laser-unpaired trials to account for changes in baseline behavior within-subject over time. Lever choice and ICSS tasks included inactive operands to account for nonspecific behavioral interactions unrelated to laser delivery.

### Statistical analysis

Males and females were included in the study but, following histological exclusion, final sample sizes were not sufficiently powered to detect sex differences so analyses were pooled. For statistical tests, we ran linear mixed-effects models using subject as a random intercept to account for variability between-subjects. We used a significance threshold (alpha) of p=0.05 for statistical annotations in figures but reported exact p-values throughout. All t-tests were two-sided. All post-hoc statistical test p-values were corrected for multiple comparisons using the Sidak method. Statistical tests were performed in R using lmerTest and emmeans packages. For the optogenetics experiments, VP→VTA and VP→mdThal groups were analyzed independently except for direct comparisons in the lever choice task and ICSS.

### Viral approach

To target VP→VTA and VP→mdThal somas in a projection-specific manner, an intersectional dual-virus strategy was used(Tervo et al., 2016). In each experiment, an AAV containing a Cre-dependent construct was injected into the VP and a retro-AAV expressing Cre recombinase was injected into the projection target region (VTA or mdThal), limiting cre-dependent expression specifically to VP neurons with terminals in the target region. GCaMP6f, an optical calcium sensor, was used as an indicator of neuronal activity in fiber photometry experiments. In optogenetic experiments, channelrhodopsin (ChR2), a light-sensitive cation channel was used for photostimulation.

### Surgery

At least one week after arrival, rats were anesthetized using 4% isoflurane for stereotaxic surgery. Rats received preoperative subcutaneous injections of carprofen (5 mg/kg) for analgesia and cefazolin (75 mg/kg) to prevent infection. An incision was made to expose the skull, and craniotomies were drilled over the VP and either the VTA or mdThal. Viral injections for VP were aimed at 0.3 mm anteroposterior (AP), +/-2.3 mm mediolateral (ML), and -8.3mm anteroposterior (AP) relative to bregma. Viral injections for VTA were aimed at mm -5.8 AP, +/-0.7 mm ML and -8.0 mm DV relative to bregma. To target VP terminals in mdThal, while avoiding terminals in the lateral habenula, viral injections were aimed at -2.5 mm AP, 0 mm ML, and -6.0 mm DV. Rats intended for fiber photometry experiments (n=11 females) received unilateral infusions of AAV9 FLEX-GCaMP6f (0.4uL, titer ∼1.4 × 10^13^ GC/mL, Addgene #100833) into the VP and retro-AAV Cre (0.4uL, titer ∼ 1.03 × 10^14^ GC/mL, Addgene #51507,) into the ipsilateral VTA at a rate of 0.1uL/min. Rats intended for optogenetic experiments received AAV9 DO-hChR2-mCherry (0.4uL, titer ∼ 3.1 × 10^13^ GC/m, Addgene # 20297) injected unilaterally into the VP and retro-AAV Cre (titer ∼ 1.1 × 10^13^ GC/mL, Addgene #51507) injected into either the ipsilateral VTA (0.4uL, VP→VTA ChR2 group; n=17, 8 females) or into the mdThal along the midline as indicated above (1.0uL, VP→mdThal ChR2 group; n=22, 10 females) at a rate of 0.1uL/min. Optical fiber cannulae (0.48 NA, 400 micron from Doric for fiber photometry; 0.29 NA, 300 micron made in-house for optogenetics) were implanted over the VP of these rats and secured by four screws to the skull with dental cement (Jet). After each surgery, rats were monitored to ensure proper recovery, receiving at least three additional daily carprofen and cefazolin injections. Optogenetic manipulation rats recovered in their home cages at least one week post-surgery prior to additional handling, at least two weeks prior to behavioral training, and at least four weeks prior to optogenetic test sessions. Fiber photometry group rats recovered in their home cages at least one week post-surgery prior to additional handling and at least four weeks prior to behavioral training to allow sufficient time for GCaMP6f expression prior to recordings.

### Discriminative stimulus task (DS task)

Rats underwent conditioning to learn a discriminative stimulus task within an operant chamber (Med Associates) as previously described (Nicola et al., 2004; Richard et al., 2016, 2018). At random intervals, an auditory discriminative stimulus (DS) was presented, signaling availability of reward (0.2mL 10% sucrose solution), contingent on port entry. To control for generalized effects of unexpected auditory stimuli, a neutral stimulus (NS) was also presented at semi-random intervals. The NS was never predictive of reward. To control for intrinsic motivational value of these auditory stimuli, the identities of the DS and NS (21khz siren noise or white noise) was counterbalanced between subjects. To promote quick and selective responding to the DS, training occurred in stages. Rats were first trained with the DS alone, which was presented for up to 60s during the first training sessions (Stage 1), and then reduced to 30s (Stage 2), 20s (Stage 3) and then 10s in subsequent training sessions (Stage 4), as long at the rats maintained responding of at least 60%. Once rats responded at least 60% of the time to the 10s DS cue, the NS was introduced (Stage 5). Rats were then trained on the full version of the DS task until they reached performance criteria of port-entry on 60% of DS trials and only 40% of NS trials. After reaching criteria, a 0.5s (Stage 6) and finally a 1.0s (Stage 7) delay was added between port-entry and reward delivery in rats tested with fiber photometry to better dissociate photometry signals related to reward anticipation and consumption.

### *In vivo* calcium recordings

#### Fiber photometry recordings and preprocessing

Rats were tethered to a fiber photometry system during training sessions by connecting their fiber optic cannulae to 300um core fiber optic patch cables (Doric) using bronze mating sleeves (Doric). The fiber photometry system consisted of excitation LEDs (465nm, 405nm; Doric), LED driver (ThorLabs) a filter cube (400-410nm isosbestic excitation filter; 460-490nm excitation filter; 500-550nm emission filter; Doric), amplifier (Doric), photometer (Doric; Newport), and data acquisition system (Tucker Davis Technologies RZ5P). Excitation LEDs were frequency-modulated (531 Hz for 465nm; 211 Hz for 405nm) and driven such that power measured at the fiber tip was ∼30uW. Isosbestic and calcium-dependent signals were demodulated as previously described (Zalocusky et al., 2016). LEDs were controlled and photometry signals were recorded using the Synapse software (Tucker Davis Technologies). Timestamps of behavioral events were also collected using TTL pulses from Med-PC (Med Associates). A 6^th^-order low-pass filter at 3Hz was applied to photometry signals at the time of collection. Following data acquisition, photometry signals were downsampled to 40Hz using custom Matlab code. Baseline correction was done independently for the raw 465nm and 405nm signals using the adaptive iteratively reweighted Penalized Least Squares (airPLS) algorithm (Zhang et al., 2010; Martianova et al., 2019). The baseline-corrected 405nm was then linearly fit to the baseline-corrected 465nm signal as previously described (Zalocusky et al., 2016). This fitted 405nm reference signal was then subtracted from the baseline-corrected 465nm to remove motion artifacts, yielding the final GCaMP6f signal. For trial-based analyses, recording sessions were divided into trials (n=30 per cue type per session), with each cue onset initiating a new trial. Photometry signals were normalized by calculating a moving Z-score on a trial-by-trial basis, using the mean and standard deviation of signal 10s prior to each cue onset as a baseline.

### Fiber photometry analyses

#### Quantifying post-cue calcium dynamics throughout training

For each trial, we computed Z-scored GCaMP traces time-locked to cue onset. To quantitatively compare magnitude of traces, we computed an area-under-the-curve (AUC) of the GCaMP trace from event onset until 10s post-event onset. To examine the relationship between learning and VP→VTA activity, we examined a subset of four specific sessions for each subject: The first day of training, the first day of Stage 5 (when the NS is introduced), the day behavioral criteria is met for Stage 5, and the final day of Stage 7. To test the impact of cue type on VP→VTA activity throughout training, we ran a linear mixed-effects model using GCaMP AUC as the response variable with fixed effects of cue type and session along with a random intercept of subject. To test if the post-cue GCaMP AUCs were nonzero, we ran post-hoc one-sample T-tests comparing each post-cue GCaMP AUC to a null distribution centered at zero. To measure changes in the magnitude of post-cue VP→VTA activity across training, we ran posthoc pairwise T-tests between sessions for each cue-type. To determine if post-DS VP→VTA activity was contingent on reward seeking behavior, we assigned each DS trial a binary port entry outcome variable which reflected whether the animal made a port entry and earned a reward. We then ran a linear mixed-effects model using post-DS GCaMP AUC as the response variable with fixed effect of port entry outcome and random intercept of subject. To test if the post-cue GCaMP AUCs were nonzero, we ran post-hoc one-sample T-tests comparing each post-cue GCaMP AUC to a null distribution centered at zero.

#### Isolating event-related calcium dynamics

For each trial, we computed Z-scored GCaMP traces time-locked to task events recorded during the trial (cue onset and first port-entry). To visualize the temporal relationship between GCaMP dynamics and event timings on a trial-by-trial basis, we constructed peri-event heatplots of individual trials’ GCaMP activity and sorted trials by latency to port-entry. To evaluate the relationship between VP→VTA bulk calcium activity and DS task events, we ran an encoding model to generate event correlation ‘kernels’, or estimates of the temporal association between each event and the GCaMP signal. To dissociate the degree to which VP→VTA activity was related to cue presentation and/or animals’ reward seeking and consumption, we included DS onset and first port-entry events in our model. The encoding model analysis only included trials from Stage 7, which had a 1.0s delay between port-entry and sucrose delivery. Trials without recorded port-entries were not included in the model. Regression kernels for each behavioral event were computed using a linear encoding model as previously described in detail(Parker et al., 2016, 2019). Briefly, timestamps for each event type during each trial were binary coded. These event timestamps were used as fixed effects in a linear regression model, where the bulk photometry signal was equal to the linear sum of fluorescence evoked by each event. Timestamps for each event were shifted by one time bin. The regression was recomputed using these shifted event timestamps. This was repeated such that each possible event time throughout the trial’s duration was represented, generating a continuous kernel of regression coefficients for each event type. AUCs of each kernel were computed from 0s to +5s post-event. To statistically compare event kernels, we ran a linear mixed-effects model using kernel AUC as the response variable with fixed effects of event type along with a random intercept of subject. To test if the kernel AUCs were nonzero, we ran post-hoc one-sample T-tests comparing each kernel AUC to a null distribution centered at zero. To evaluate the encoding model’s accuracy, we ran a correlation between the true peri-DS GCaMP signal and the model’s predicted peri-DS GCaMP signal generated by event timings.

#### Latency correlation analysis

Correlation coefficients between port-entry latency and photometry signals were computed on a trial-by-trial basis for each timestamp. Briefly, the photometry signals for each trial were time-locked to cue onset and converted to a Z-score using 10s preceding cue onset as a baseline. Photometry signals following the first port entry timestamp during each trial were excluded to eliminate contamination from consumption-related fluorescence. The latency correlation analysis only included trials from Stage 7, which had a 1.0s delay between port-entry and sucrose delivery. Pearson correlations were then computed between the photometry signal and port-entry latency for each timestamp in the peri-cue time window. Correlation coefficients were averaged between all trials for each subject, then averaged between subjects to yield between-subject mean coefficients. To compare resulting coefficients against chance, we also ran a shuffled version of this correlation analysis in which port-entry latencies were shuffled randomly between trials prior to correlation and ran a linear mixed-effects model using coefficient as the response variable with fixed effects of shuffle and time bin along with a random intercept of subject.

### *In-vivo* optogenetic manipulations

#### Optogenetic excitation during the DS task

Rats tested for the effects of optogenetic stimulation in the DS task were trained while connected via ceramic mating sleeves to 200 µm core patch cords, which were connected to a 473 nm DPSS laser via a fiber optic rotary joint mounted on a cantilever arm. After rats met training criteria in the DS task, we assessed the impact of activating VP→VTA neurons during DS presentations. To ensure rats discriminated between cues prior to optogenetic stimulation, rats were only included in analyses if they met performance criteria on the day prior to test day 1 (pre-test). Criteria was met when a rat’s port-entry probability on DS trials was least 60% and at least twice their port-entry probability on NS trials. On test day 1, rats received 1s of 20Hz stimulation (465 nm laser, 10-15 mW) during 50% of DS and NS presentations (randomly selected) at the start of the cue. On test day 2, rats received up to 10s of 20 Hz stimulation on 50% of cue presentations; this stimulation co-terminated with port-entry during the cue. We assessed the impact of this stimulation on port entry latency and probability. We ran linear mixed-effects models for each test day (pre-test, 1s, or >=10s stimulation) using port entry probability or port entry latency as the response variable with fixed effects of cue type and laser delivery with a random intercept of subject.

#### Choice task with optogenetic excitation

To assess whether activation of VP→VTA neurons during reward consumption would enhance the reward’s value and bias choice behavior, we trained rats in a choice task with optogenetic stimulation. During the task, rats were presented with two levers. Presses on either lever resulted in a 2s (.2mL) delivery of 10% sucrose to the reward port and retraction of the levers. Presses on one lever (the “stimulation” lever) also resulted in optogenetic stimulation paired with reward consumption, whereas the other lever did not produce any laser stimulation. Following presses on the stimulation lever, the first lick of 10% sucrose triggered 20 Hz stimulation (465 nm, 10-15 mW), which continued for up to 8s or until the rat stopped licking for at least 0.5s. The levers were reinserted 8s after reward consumption was initiated. To assess the impact of optogenetic activation of VP→VTA neurons on reward-seeking and consumption, we assessed the total number of presses on each lever, the proportion of presses on the stimulation lever, as well as licks per reward with and without stimulation. Licks per reward were only calculated and included in analyses from sessions where rats completed at least 3 lever presses of each type to ensure that both were sampled sufficiently for comparisons. We ran a linear mixed-effects model using lever press count or licks per reward as the response variable with fixed effects of lever type and session with a random intercept of subject. We also ran a linear mixed-effects model using proportion of active lever presses as the response variable with a fixed effect of session and random intercept of subject. To compare these proportions against chance, we ran post-hoc one-sample T-tests comparing each session’s active proportion to a null distribution centered at 0.5.

#### Intracranial self-stimulation (ICSS)

To assess whether activation of the VP→VTA pathway was rewarding on its own, as reported previously (Faget et al., 2018). we assessed optogenetic self-stimulation of this pathway in five one-hour sessions. During these sessions, entry into the “active” nosepoke resulted in 1s 20 Hz stimulation (465 nm laser, 10-15 mW) and simultaneous illumination of the nosepoke, whereas entry into the “inactive” nosepoke resulted in illumination of the nosepoke, but no optogenetic stimulation. To assess the impact of optogenetic activation of the VP→VTA pathway on reward-seeking and consumption, we assessed the total number of entries into each nosepoke and the proportion of active nosepokes with and without stimulation. We ran linear mixed-effects models using nosepoke count as the response variable with fixed effects of nosepoke type and session with a random intercept of subject. We also ran a linear mixed-effects model using proportion of active nosepokes as the response variable with a fixed effect of session and random intercept of subject. To compare these proportions against chance, we ran post-hoc one-sample T-tests comparing each session’s active proportion to a null distribution centered at 0.5.

### Histology

Following the completion of behavioral testing, rats were anesthetized using pentobarbitol (390mg/mL; Fatal-Plus) and transcardially perfused with cold 1x PBS and 4% PFA. Brains were extracted and post-fixed at 4C in 4% PFA overnight before being transferred to a 20% sucrose in 1x PBS solution for cryoprotection. Brains were left in sucrose solution until they sank completely (at least 48 hours). 50um coronal sections were collected using a cryostat (Leica Biosystems). Sections were stored at 4C in 1x PBS before being mounted on glass slides using Vectashield antifade mounting medium with DAPI (Vector Laboratories). Images of sections were collected using a fluorescent microscope (Nikon TI-E Deconvolution Microscope) with a 4x objective and camera (ORCA Flash 4.0). Fiber positions and viral expression (GCaMP6f or ChR2-mCherry) within VP were confirmed by registering images with a reference atlas (Paxinos & Watson, 7^th^ Edition) in ImageJ. Two animals from the photometry group were excluded from analysis due to insufficient GCaMP expression or loss of photometry signal during the experiment. Three animals from the VP→VTA ChR2 group and six animals from the VP→mdThal ChR2 group were excluded from analysis due to insufficient ChR2 expression or fiber implant placement outside of VP.

## RESULTS

### Rats learn to discriminate between the DS and NS

To record calcium activity from VP→VTA neurons during the performance of cue-elicited reward seeking, a genetically encoded calcium indicator, GCaMP6f, was expressed using an intersectional viral approach (Figure 1A) and optical fibers were implanted in VP (Figure 1B).

**Figure 1.**
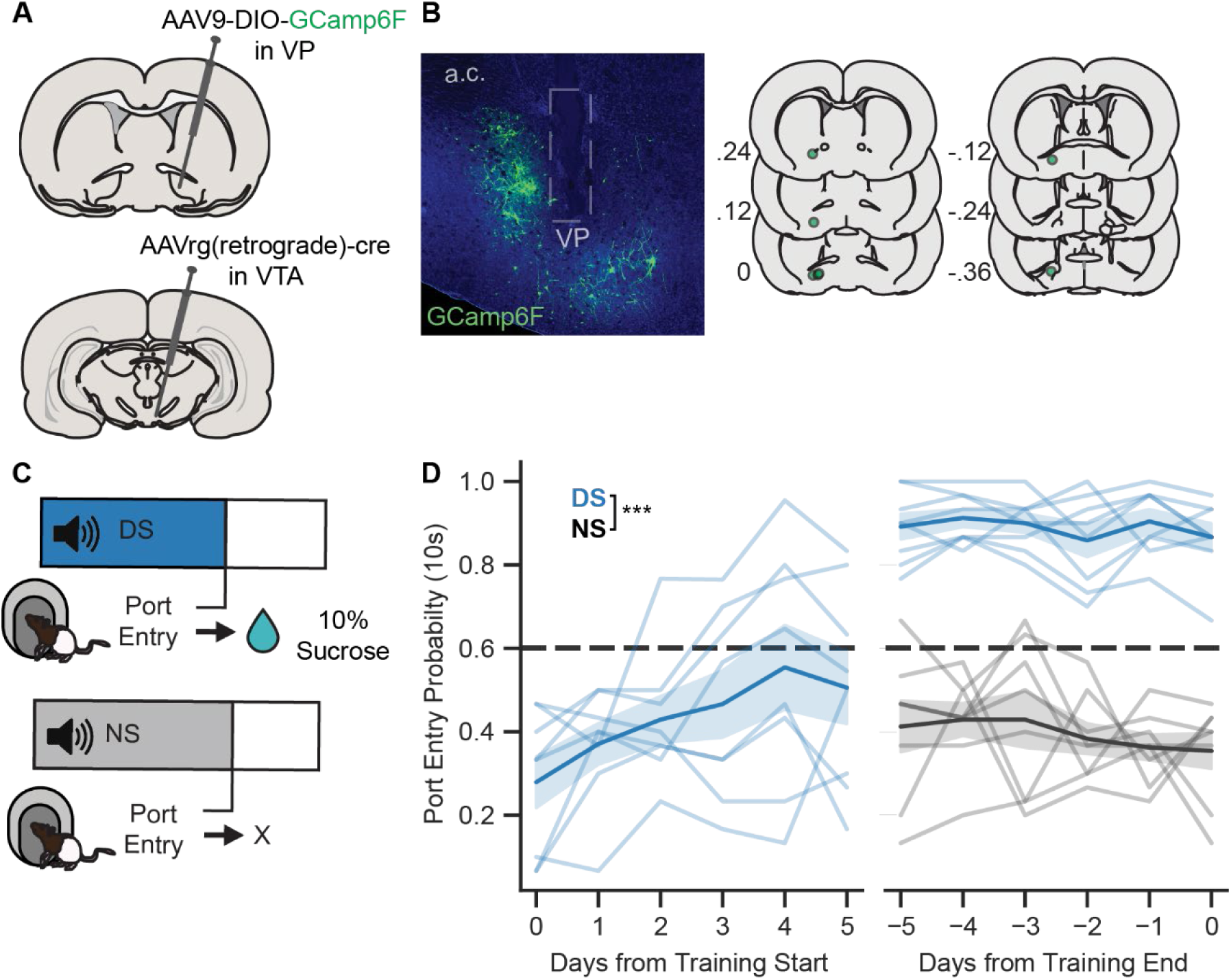
Viral approach and DS Task. A) Diagram of intersectional viral approach used for projection-specific GCaMP6f recordings from VP→VTA. B) Representative image of GCaMP6f expression (green) and optic fiber tract (gray outline) in the VP (“a.c.” denotes anterior commissure). C) Diagram of DS task design, with 10% sucrose reward delivery contingent on rats’ port-entry during DS presentation. D) Mean probability of port-entry within 10s of cue onset (blue= DS, gray= NS) in the first 5 and final 5 training sessions (main effect of cue, F(1, 77)= 534.297, p<0.001). Lines with shading indicate between-subjects mean +/-SEM (n=8 rats).

Four weeks following recovery from surgery, rats were trained to perform a discriminative stimulus task (DS task). In the DS task, presentation of an auditory reward-predictive DS signaled availability of 10% sucrose contingent on rats’ port-entry, while the NS was not associated with reward availability (Figure 1C). To assess DS task performance, we calculated the probability and latency to enter the port within 10s following cue onset for each session as previously described (Nicola et al., 2004; Richard et al., 2016). As expected, we found that the probability of port-entry within 10s of DS onset, but not NS onset, increases across sessions (Figure 1D, mean +/-SEM days to reach Stage 5 criteria = 17.375 +/-3.184). By the end of training, rats consistently responded more often to the DS than to the NS (main effect of cue, F(1, 77)= 534.297, p<0.01) In addition, rats made port-entries more quickly following DS presentations compared to NS presentations (data not shown; port-entry latency, main effect of cue, F(1,1857.9)= 196.235, p<0.001). We excluded 2 subjects from analyses due to a lack of GCaMP6f expression under the fiber implant and excluded 1 additional subject that failed to meet training criteria. These data demonstrate that rats included in subsequent analyses behaviorally discriminated between DS and NS, selectively responding to the reward-predictive cue.

### VP→VTA neurons are selectively active on DS trials, contingent on port-entry

To determine if VP→VTA calcium activity exhibited reward-related changes, we first compared the GCaMP signal between DS and NS trials (Figure 2A). To quantitatively compare the magnitude of GCaMP responses, we computed peri-cue AUCs of the GCaMP signal during the 10s following cue onset (Figure 2B). As expected, we observed that VP→VTA calcium activity was significantly elevated on DS trials compared to NS trials across training stages (main effect of cue, F(1,1427) = 122.858, p < 0.001). T-tests revealed that DS AUCs were significantly different from zero in each session (DS vs. zero; first session, t(6)= 2.48, p= 0.048; NS introduced, t(14.6)= 5.397, p<0.001; Criteria met, t(14.6)= 7.447, p<0.001; Final session, t(14.6)= 9.908, p<0.001) while the NS AUCs were only significantly different from zero in the first session the NS was present (NS vs. zero; NS introduced, t(14.6)= 2.699, p=0.050; Criteria met, t(14.6)= 2.199, p=0.127; Final session, t(14.6)= 1.356, p=0.480). The magnitude of the peri-cue GCaMP AUC increased throughout the course of training for DS trials, but not NS trials (interaction of cue type and session, F(2,1635.9) = 11.193, p < 0.001; no significant pairwise follow-up comparisons between sessions for NS). These data suggest that VP→VTA are selectively active following reward-cue presentation and do not respond to neutral cues. The increase in DS AUC magnitude across sessions may reflect animals’ learning.

**Figure 2.**
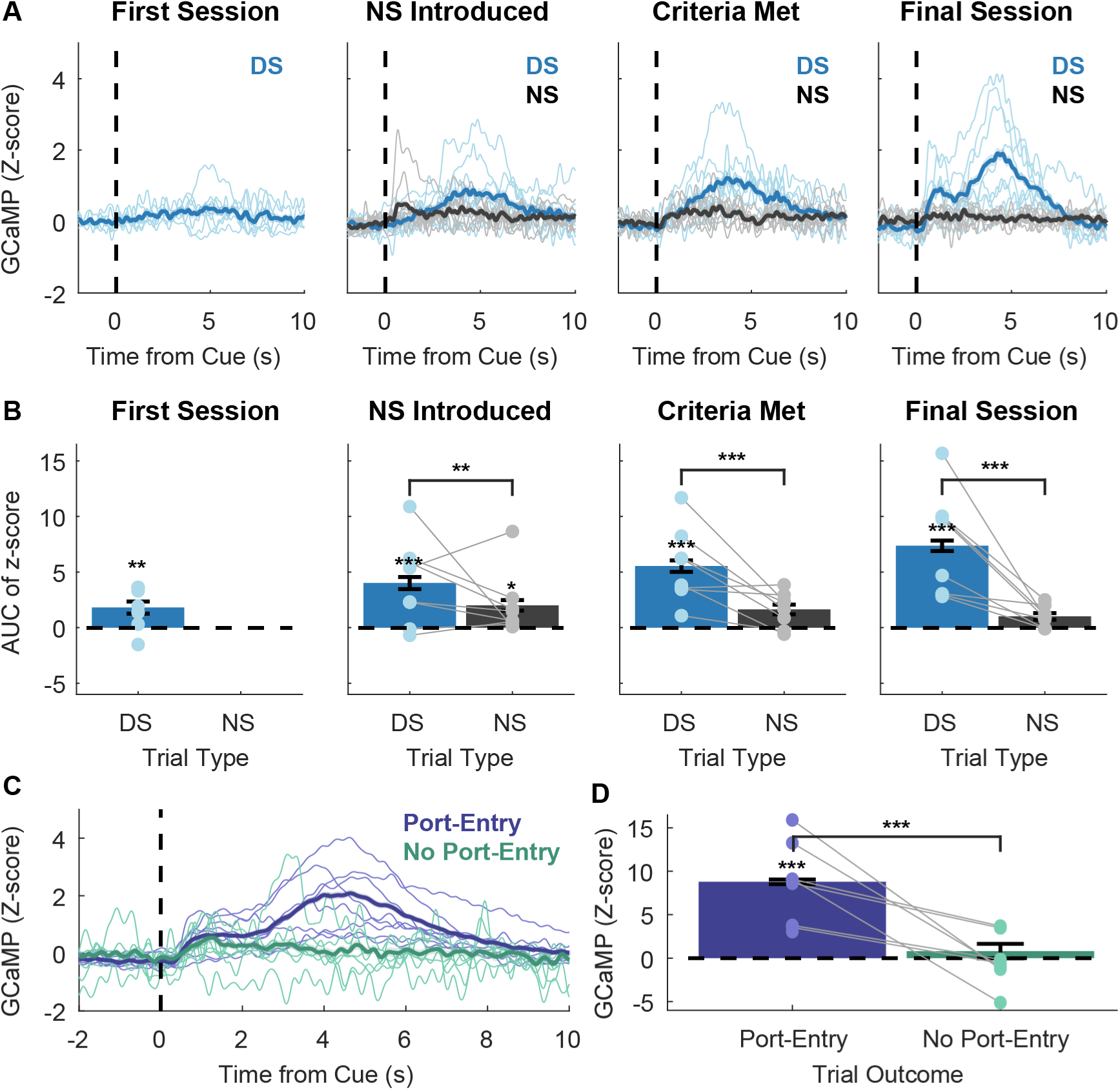
Calcium dynamics of VP→VTA during DS task performance. A) Mean peri-cue calcium traces (blue= DS, gray=NS) on day 1 of training, first day of Stage 5 (when NS is introduced), the day Stage 5 criteria was reached, and the day Stage 7 criteria was reached. Lines with shading indicate between-subjects mean +/-SEM (n=8 rats). Individual points and lines without shading indicate individual subject means. B) Mean AUC of peri-cue calcium traces on day 1 of training, the first day of Stage 5, the day Stage 5 criteria was reached, and the day Stage 7 criteria was reached (Main effect of cue (F(1,1427)= 122.858, p<0.001; DS vs. zero; first session, t(6)= 2.48, p = 0.048; NS introduced, t(14.6)= 5.397, p<0.001; Criteria met, t(14.6)= 7.447, p<0.001; Final session, t(14.6)= 9.908, p<0.001; NS vs. zero; NS introduced, t(14.6)= 2.699, p=0.050). C) Mean Stage 7 peri-DS calcium traces by behavioral outcome (purple= port-entry during DS, teal= no port-entry during DS). D) Mean AUC of Stage 7 peri-DS calcium traces by behavioral outcome (main effect of port-entry outcome F(1,1279.5)= 51.047, p<0.001; DS with port-entry vs. zero, t(7.02)= 5.217, p=0.002).

To examine whether VP→VTA activity was reflective of DS presentation alone or associated with subsequent reward-seeking behavior, we compared peri-DS GCaMP traces from trials with a port-entry outcome to those from trials without a port-entry. Interestingly, significant peri-DS VP→VTA calcium activation was present only on DS trials in which the animals entered the port (Figure 2D; DS with port-entry versus DS with no port-entry, F(1, 1279.5)= 51.047, p<0.001; DS with port-entry versus zero, t(7.02)= 5.217, p=0.002; DS with no port-entry versus zero, t(11.92)= 0.834, p=0.664). These data suggest that peri-DS VP→VTA activation may not be elicited by the DS cue itself but may be associated with subsequent pursuit and/or consumption of reward. Alternatively, the cue itself may evoke a stronger response on trials when the animal is more motivated to respond, as has been previously reported in VP (Richard et al., 2016, 2018). Therefore, our subsequent analyses aimed to more precisely examine the temporal associations between the discrete task events (cue versus port entry) and the GCaMP signal.

### Population-level VP→VTA calcium activity is dynamic and peaks during reward consumption

To examine the temporal relationship between the GCaMP photometry signal and behavioral task events, we examined peri-event traces time-locked to distinct task events (DS onset, first port-entry following DS; Figure 3A). Qualitatively, two peaks were apparent in the peri-DS traces: the first with an initial rise in signal starting from +0.3s to a peak centered around +1.0s, followed by the second with a greater rise starting from +2.5s to a peak centered around +4.0s. In the traces time-locked to port-entry, these two peaks were also present but were greater in magnitude and are shifted earlier in time relative to peri-DS traces, revealing a temporal dependence on port-entry. Interestingly, when time-locked to port-entry, the initial ramp in GCaMP signal appeared to begin prior to port-entry. To better examine the calcium activity in response to DS versus port-entry in the context of trial-to-trial variability in response latency, we made peri-event heat plots for individual trials, time-locked to the DS versus port-entry and sorted based on the latency between task events for each trial (Figure 3B, representative subject). Peri-event heat plots revealed that the second increase in VP→VTA GCaMP fluorescence consistently occurs after port entry, shifting later in time as response latency grows longer, peaking during sucrose consumption (post-lick). These heat plots revealed variable dynamics in the epoch between cue onset and port-entry which may reflect responses to cue presentation and/or signals related to reward pursuit. Therefore, we proceeded with a linear encoding model analysis to statistically evaluate the relationship between event timing and calcium activity (Parker et al., 2016, 2019).

**Figure 3.**
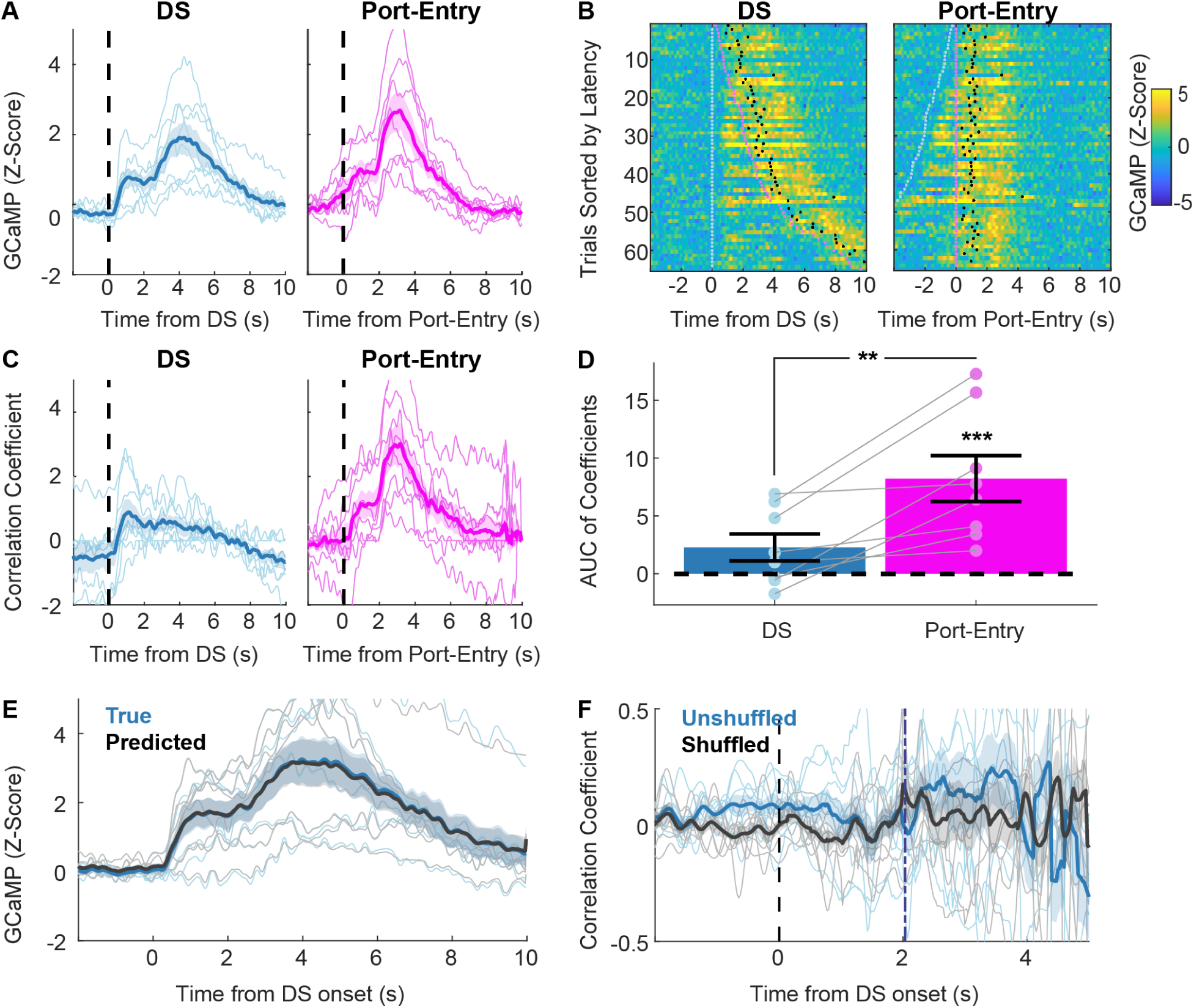
Temporal relationship between DS trial events and VP→VTA calcium activity. A) Mean Stage 7 peri-event calcium traces time-locked to distinct task events (teal= DS onset, purple= first port-entry). Lines with shading indicate between-subjects mean +/-SEM (n=8 rats). Individual points and lines without shading indicate individual subject means. B) Peri-event heat plots of GCaMP signal from Stage 7 DS trials with port-entry outcome (n=1 representative subject). Trials sorted by port-entry latency with event onset points overlaid (teal= DS onset, purple= first port-entry, magenta= first lick). C) Mean regression kernel traces for DS trial events (teal= DS onset, purple= first port entry) generated by encoding model. Lines with shading indicate between-subjects mean +/-SEM (n=8 rats). Individual points and lines without shading indicate individual subject kernels. D) Distribution of event kernel AUCs from 0s and +5s (main effect of event type F(1,7)= 14.372, p=0.007; Port-entry vs. zero, t(10.9)= 5.053, p<0.001). Individual points and lines without shading indicate individual subject AUCs. D) Mean true (blue) and predicted (black) peri-DS onset GCaMP traces from encoding model trials (*r*(3838)= 0.997, p<0.001). Lines with shading indicate between-subjects mean +/-SEM (n=8 rats). Individual points and lines without shading indicate individual subject means. E) Correlation between GCaMP signal at each time bin sampled and port-entry latency (blue= true latency, gray= shuffled latency). Purple dashed line represents mean port-entry latency. Lines with shading indicate between-subjects mean +/-SEM (n=8 rats). Individual points and lines without shading indicate individual subject means.

To evaluate the relationship between VP→VTA bulk calcium activity and DS task events, we ran an encoding model to generate event correlation “kernels’’ or estimates of the temporal association between each event and the GCaMP signal (Figure 3C). To dissociate the degree to which VP→VTA activity was related to cue presentation and/or animals’ reward seeking, we included DS onset and first port-entry events in our model. We did run a version of the encoding model which included the first lick event as well, but found that first lick events were highly collinear with the first port-entry events and thus did not improve the model’s fit (data not shown), so we proceeded with the simpler two-event model. Briefly, the encoding model used shifted versions of the event onset timings (with one shift for every possible time bin in a trial from -2s-10s post-DS onset) as predictors in a regression. Thus, resulting regression kernels represent the best fit coefficients, or predictive value, of each event throughout the time course of a trial (Figure 3C). To quantitatively compare event kernels, we computed AUCs from 0-5s (Figure 3D). We found that the AUC of the port-entry kernel, but not the DS kernel, was significantly different from zero (t(10.9)= 5.053, p<0.001). In addition, port-entry kernel AUCs were greater in magnitude than the DS kernel AUC (main effect of event type F(1,7)= 14.372, p=0.007; pairwise difference t(7)= 3.791, p=0.007). Importantly, the model accurately predicted the actual GCaMP signal for each trial using event timings alone (Figure 3E; *r(*3838)= 0.997, p<0.001). The encoding model results suggest that VP→VTA activity on DS trials is not driven by cue presentation, but is primarily related to events following port-entry, presumably reward delivery and consumption.

### Population-level VP→VTA calcium activity does not encode the vigor of cue-elicited reward-seeking

Prior extracellular recordings identified a subset of VP neurons which encode incentive motivational value of cues during the DS task, measured by port-entry latency (Richard et al., 2016, 2018). We have also found similar correlations between port-entry latency and peri-cue population-level calcium activity in VP GABA neurons (Scott et al., 2022). To determine whether VP→VTA calcium activity on DS trials is predictive of reward-seeking vigor, we ran a correlation between the peri-cue, pre-port-entry GCaMP6f signal in each 0.025s time bin sampled and latency of port-entry on a trial-by-trial basis (Figure 3F). To compare resulting coefficients against chance, we also ran a shuffled version of this correlation analysis in which port-entry latencies were shuffled randomly between trials prior to correlation. Contrary to our initial hypothesis, we found no significant correlation between VP→VTA activity and port-entry latency, or pairwise differences between the true and shuffled correlations (true versus shuffled, F(1,6511) =59.910, p <0.001 ; time, F(575,6511) = 1.256, p < 0.001).

### Acute cue-paired optogenetic stimulation of VP→VTA neurons does not impact probability or vigor of reward-seeking

To determine whether VP→VTA causally promotes cue-elicited reward seeking, we performed cue-paired optogenetic manipulations of this pathway during the DS task by virally expressing ChR2 and delivering laser through fiber optic implants (Figure 4A). We tested the impact of VP→VTA excitation in two separate sessions, defined by maximum laser duration (Figure 4B). To emulate transient “cue” responses reported in prior recordings from VP neurons, we ran 1s stimulation sessions where 50% of cues were paired with intracranial laser delivery (20Hz), starting at cue onset and terminating after 1s. To manipulate VP activity throughout the entire cue duration, we ran <=10s stimulation sessions where 50% of cues were paired with intracranial laser delivery (20Hz), starting at cue onset and terminating once the animal made a port-entry or at 10s if the animal failed to enter the port during the cue. To determine the specificity of behavioral effects to VP→VTA neurons, we performed the same manipulations in another VP output pathway, VP→mdThal.

**Figure 4.**
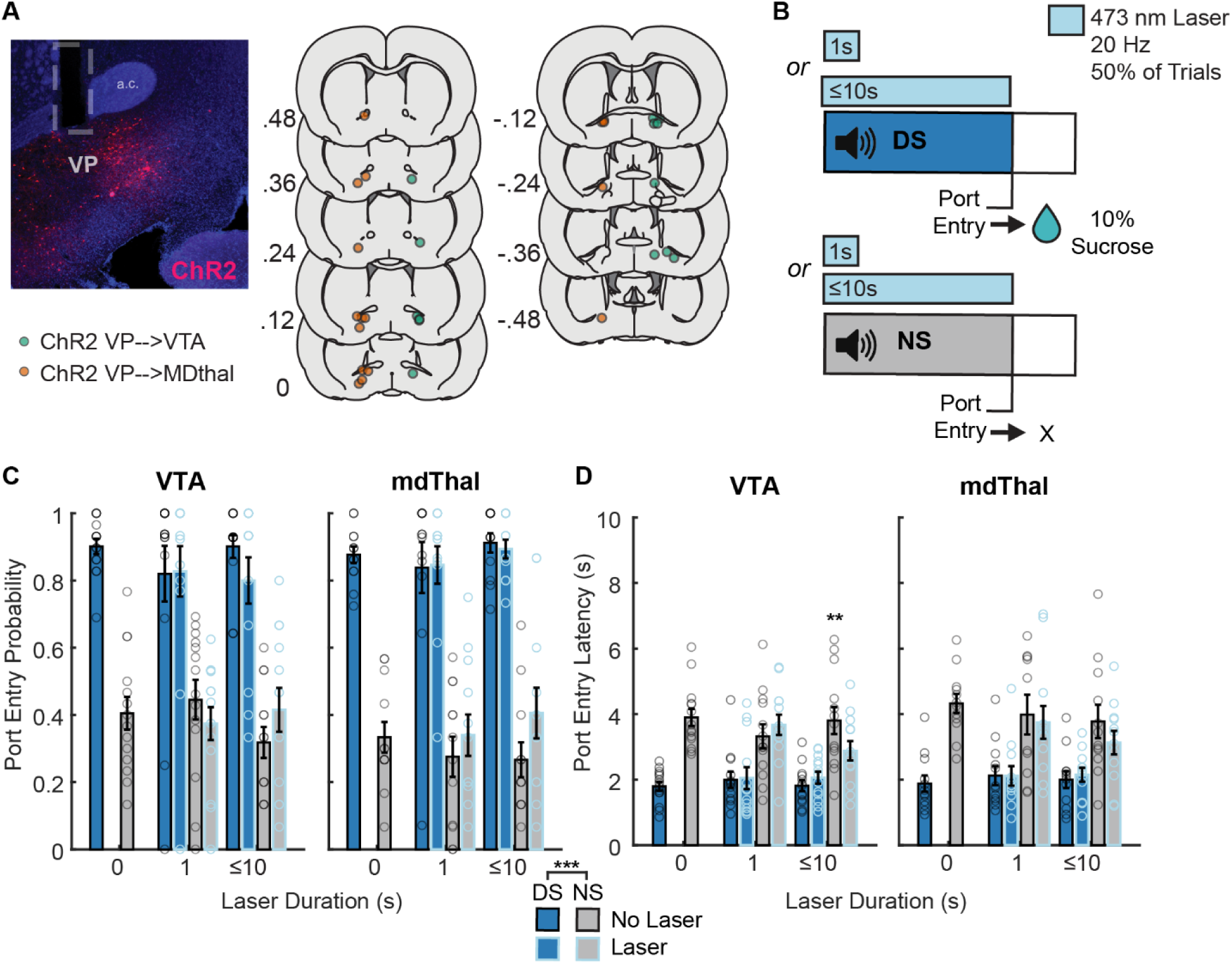
Cue-paired optogenetic stimulation of VP→VTA and VP→mdThal neurons during the DS task. A) Left-Representative image of ChR2 expression (red) and optic fiber tract (gray outline) in the VP (“a.c.” denotes anterior commissure), Right-Coordinates of optic fiber implant tips of subjects in VP→VTA ChR2 group (green) and VP→MDthal ChR2 group (orange). B) Diagram of DS task sessions with cue-paired optogenetic stimulation, where 50% of cue presentations (DS=blue, NS=gray) are paired with either 1s (acute 1s stimulation sessions) or <=10s (<=10s cue-paired sessions) intracranial laser delivery (473nm, 20Hz). C) Mean probability of port-entry of VP→VTA ChR2 group (Left) and VP→mdThal group (Right) on trials (blue= DS, gray= NS) with and without laser (black outline= no laser, blue outline= laser-paired) on sessions prior to optogenetic stimulation, 1s acute stimulation sessions, and <=10s cue-paired sessions. Bars and errorbars indicate between-subjects mean +/-SEM. Individual points indicate individual subject means. D) Mean latency to port-entry of VP→VTA ChR2 group (Left) and VP→mdThal group (Right) on trials (blue= DS, gray= NS) with and without laser (black outline= no laser, blue outline= laser-paired) on sessions prior to optogenetic stimulation, 1s acute stimulation sessions, and <=10s cue-paired sessions. Bars and error bars indicate between-subjects mean +/-SEM. Individual points indicate individual subject means (VP→VTA latency to port-entry; cue × laser interaction, F(1,36) = 6.034, p = 0.0190; <=10s stimulation laser-paired NS trials vs. laser-unpaired NS trials, t(36) = 2.759, p = 0.0185).

Prior to optogenetic manipulation sessions, rats in both groups learned to preferentially respond to reward cues. This was reflected in both an increased probability of port-entry during the DS (VP→VTA main effect of cue F(1,26) = 85.438, p < 0.001; VP→mdThal, main effect of cue, F(1,22) = 108.96, p < 0.001) and faster port entries during the DS than the NS (VP→VTA main effect of cue F(1,13)= 59.756, p < 0.001; VP→mdThal, main effect of cue F(1,11)= 76.163, p<0.001). When we tested the impact of optogenetic excitation on 50% of trials, we found no significant impact of acute 1s optogenetic stimulation of VP→VTA on port-entry probability (Figure 4C; main effect of laser F(1,39)= 0.479, p=0.493; interaction of cue and laser F(1,39)= 0.708, p=0.405) or latency (Figure 4D; effect of laser F(1,36)= 0.545, p=0.465; interaction of cue and laser F(1,36)= 0.298, p=0.589). We also found no impact of acute 1s optogenetic stimulation of VP→mdThal on port-entry probability (main effect of laser F(1,33)= 0.519, p=0.476; interaction of cue and laser F(1,33)= 0.320, p=0.575) or latency (main effect of laser F(1,30.215)= 0.274, p=0.605, interaction of cue and laser F(1,30.215)= 0.246, p=0.623). In these 1s optogenetic manipulation sessions, rats in both groups maintained a significantly higher probability of responding to the DS than the NS (VP→VTA, main effect of cue, F(1,39) = 79.116, p < 0.001; VP→mdThal, main effect of cue, F(1,33)= 116.289, p<0.001). Both groups also continued to enter the port more quickly in response to the DS than the NS (VP→VTA, main effect of cue, F(1,36) = 30.272, p < 0.005; VP→mdThal, main effect of cue, F(1,30.215) = 22.565, p < 0.001). These results suggest that activation of either pathway on the timescale on which correlations between VP single unit activity and latency have been reported is insufficient to alter the likelihood or latency of cue-elicited reward seeking.

In the <=10s optogenetic stimulation sessions, we found no significant effect of optogenetic stimulation in the VP→mdThal group on either port-entry probability (Figure 4C; main effect of laser, F(1,33)= 1.639, p=0.209; interaction of cue and laser F(1,33)= 2.782, p=0.105) or latency (Figure 4D; main effect of laser, F(1,33)= 0.701, p=0.409; interaction of cue and laser, F(1,33)= 1.839, p=0.184). These animals maintained preferential responding to the DS with an increased port-entry probability (main effect of cue, F(1,33)= 144.680, p<0.001) and shorter port-entry latency (main effect of cue, F(1,33)=22.171, p<0.001). In contrast, optogenetic stimulation <=10s altered cue responses in the VP→VTA group, especially on NS trials. Specifically, activation of VP→VTA impacted port-entry latency in a cue-specific manner (interaction of cue and laser, F(1,36) = 6.034, p = 0.0190), in that rats entered the port more quickly on NS-paired laser trials compared to NS trials without laser (t(36) = 2.759, p = 0.0185). While we observed a significant cue by laser interaction for port-entry probability (F(1,36) = 4.204, p = 0.0477), this was not explained by any significant post-hoc pairwise differences between laser and no lasers trials for either the DS or the NS, though port-entry probability appeared slightly lower on average on DS laser trials, and slightly higher on average on NS laser trials, suggested some disruption in discrimination. Overall, we did not see any significant effects of pathway-specific excitation on DS-evoked behavior.

### Sucrose-paired optogenetic stimulation of VP→VTA neurons, but not VP→mdThal neurons, biases choice

To determine the impact of reward-paired VP→VTA or VP→mdThal neuron activation on instrumental reward seeking and consumption, rats in both groups performed a lever choice task with optogenetic stimulation (Figure 5A). In this task, an “inactive” lever resulted in sucrose delivery alone while an “active” lever resulted in sucrose delivery along with lick-paired optogenetic stimulation.

**Figure 5.**
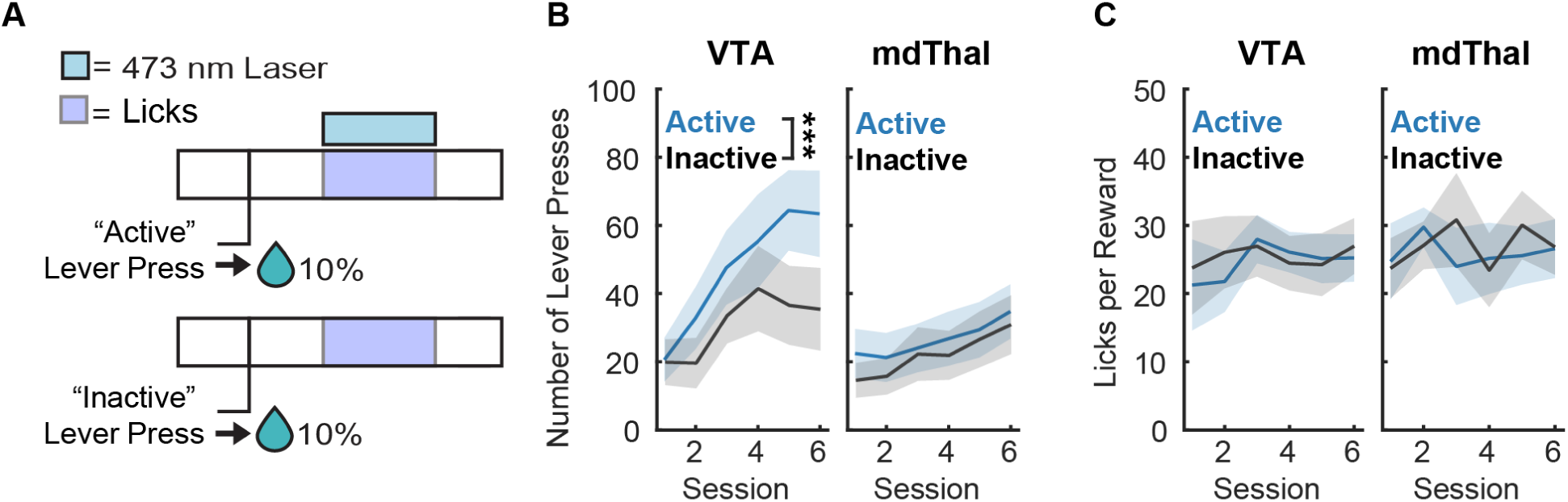
Lever choice task with reward-paired optogenetic stimulation. A) Diagram of lever choice task design, in which an “inactive” lever resulted in sucrose delivery alone while an “active” lever resulted in sucrose delivery along with lick-paired optogenetic stimulation. B) Mean number of lever presses (blue= active, gray= inactive) during lever choice task sessions for VP→VTA ChR2 group (Left; main effect of lever F(1,154)= 8.132, p=0.005) and VP→mdThal ChR2 group (Right). Lines with shading indicate between-subjects mean +/-SEM. C) Mean licks per reward delivered during lever choice task sessions for the VP→VTA ChR2 group (Left) and VP→mdThal ChR2 group (Right).

We found that the VP→VTA, but not the VP→mdThal group, developed a preference for the laser-paired lever (Fig. 5B), measured by raw lever press count (VP→VTA, main effect of lever, F(1,154)= 8.132, p=0.005; VP→mdThal, main effect of lever, F(1,165)= 1.684, p=0.196). It is possible that VP→VTA rats developed an active lever preference because optogeentic stimulation of VP→VTA neurons enhanced the hedonic value of sucrose. Since frequency of licking is associated with a substance’s hedonic value in rodents (Spector and St. John, 1998; Wassum et al., 2009; Katsuura et al., 2011; Dwyer, 2012; Riordan and Dwyer, 2019), we compared licks per reward delivered with or without lick-paired laser delivery to determine this. We found no significant impact of lick-paired optogenetic stimulation on licking behavior during these sessions (Figure 5C; VP→VTA, main effect of lever type, F(1,82.155)= 0.290, p=0.592, interaction of lever type and session, F(5,82.155)= 0.393, p=0.852; VP→mdThal, main effect of lever type, F(1,84.05)= 0.1689, p=0.682, interaction of lever type and session, F(5,84.050)= 0.3743, p=0.865), suggesting that preference for the active lever in the VP→VTA group did not depend on any impact on hedonic evaluation of the sucrose itself during stimulation. Since optogenetic stimulation in this task required execution of a complex action sequence (active lever press→reward cup entry→ licking), we ran ICSS sessions to directly test if optogenetic stimulation of VP→VTA or VP→mdThal can reinforce a new operant response.

### Stimulation of VP→VTA neurons is reinforcing

Optogenetic manipulations in the DS task and choice task occurred concurrently with cue and reward consumption, respectively. To assess the sufficiency of VP→VTA neurons to reinforce behavior in the absence of primary reward and associated cues, rats completed ICSS sessions in which an “active” nosepoke on one side of the chamber yielded optogenetic stimulation (Fig. 6).

**Figure 6.**
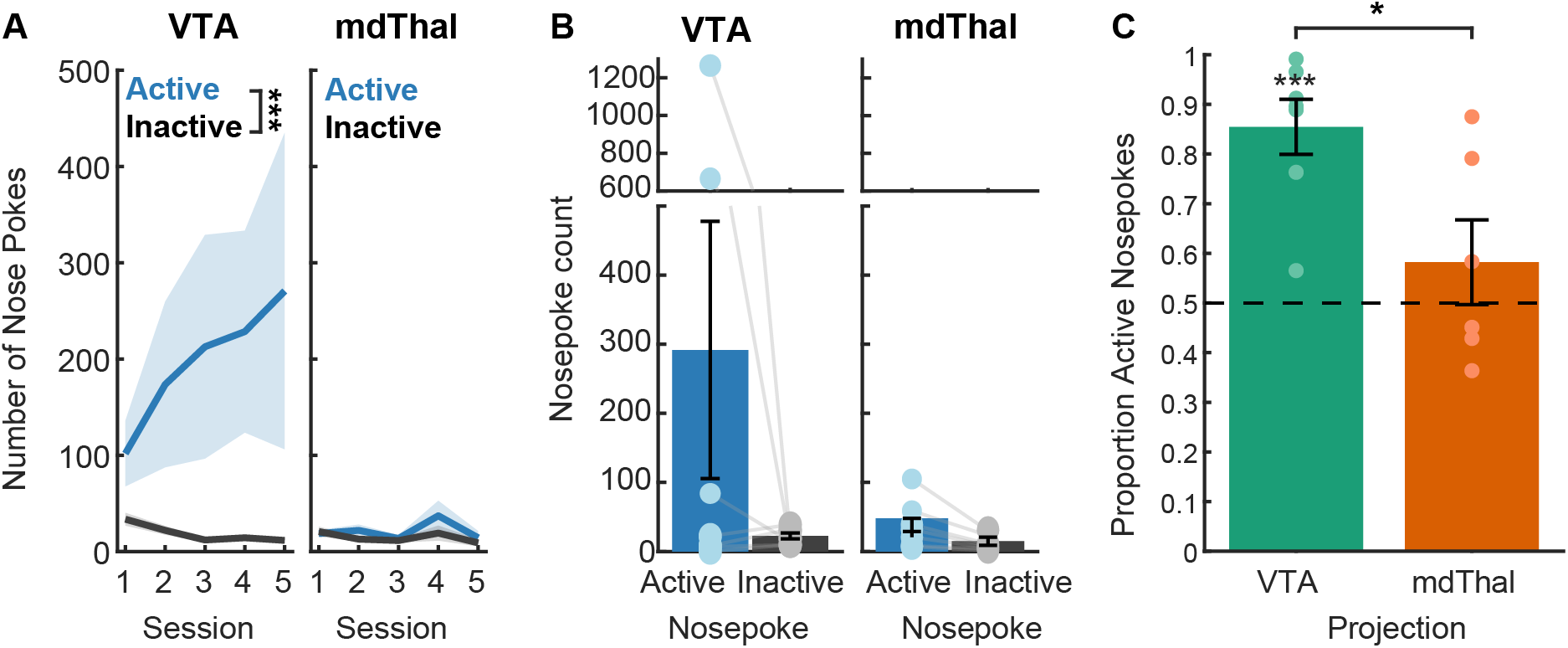
Intracranial self-stimulation (ICSS) task. A) Mean number of nose pokes (blue= active, gray= inactive) for VP→VTA ChR2 (Left; main effect of nose poke F(1,54)= 20.248, p<0.001) and VP→mdThal (Right; effect of nose poke F(1,54)= 3.505, p= 0.066) groups in the initial training phase. Lines with shading indicate between-subjects mean +/-SEM (n=). B) Mean number of nose pokes (blue= active; gray= inactive) for VP→VTA ChR2 (Left; effect of nosepoke F(1,5.974)= 2.484, p=0.166) and VP→mdThal ChR2 (Right; effect of nosepoke F(1,6)= 0.836, p=0.396). Bars indicate between-subjects mean +/-SEM. Individual points and lines indicate individual subject means. C) Mean proportion of active nosepokes for each viral group (green= VP→VTA ChR2, significant difference from chance t(6)= 6.393, p<0.001; orange= VP→mdThal)

In these ICSS sessions, VP→VTA group rats preferentially entered the active nosepoke, measured by nosepoke count (Fig 6A; main effect of nosepoke type, F(1,54)= 20.248, p<0.001; interaction of nosepoke type and session, F(4,54)= 0.669, p= 0.616) and proportion of active nosepokes significantly different from chance in the final session (Fig 6C; t(6)= 6.393, p<0.001). These data are consistent with other reports that optogenetic stimulation of VP→VTA is reinforcing (Faget et al., 2018). In contrast, VP→mdThal group pressed active and inactive nosepokes at similar rates, failing to engage in significant self-stimulation (main effect of nosepoke type, F(1,54)= 3.505, p=0.067, interaction of nosepoke type and session F(4,54)= 0.958, p=0.438). In the final session, the VP→mdThal group’s proportion of active nosepokes did not differ from chance (t(5)= 0.963, p=0.380), and was significantly lower compared to the VP→VTA group (effect of projection F(1)= 7.601, p=0.019). Overall, the ICSS data suggest that VP outputs to the VTA, but not the mdThal, are powerfully and immediately reinforcing.

## DISCUSSION

In this study, we used fiber photometry to record calcium activity of VP neurons projecting to the VTA in rats while they performed cue-elicited reward seeking (DS task). We found VP→VTA calcium responses on DS trials signaling reward availability, but not NS control trials. Using an encoding model to quantify the temporal relationship between calcium activity and task events, we report that population-level VP→VTA calcium activity primarily increases following port-entry. Critically, we did not find a significant correlation between calcium activity and response latency. We also used *in-*vivo optogenetics to acutely manipulate activity of VP→VTA neurons as well as a distinct VP output pathway, VP→mdThal, during cue presentations in the DS task. We found no impact of DS-paired VP→VTA stimulation on probability or latency of port-entry. In addition, we found that optogenetic stimulation of VP→VTA neurons paired with sucrose-consumption biases choice in a lever-choice task and that VP→VTA optogenetic stimulation alone is sufficiently reinforcing to support ICSS. Overall, these data suggest that VP→VTA neurons are robustly recruited during sucrose consumption and that these neurons support reinforcement but do not encode the vigor of reward-seeking or the incentive motivational value of cues.

### Lack of incentive value and vigor encoding by VP→VTA neurons in the DS task

Though prior work has reported vigor encoding by single-units within VP (Richard et al., 2016, 2018; Lederman et al., 2021) and population-level recordings from GABAergic VP neurons (Scott et al., 2022) in the DS task, we did not find vigor encoding in population-level VP→VTA calcium recordings. In addition, DS-paired optogenetic stimulation of VP→VTA neurons did not change response vigor. These results suggest that activation of these neurons at the timescale on which correlations between VP single unit activity and latency have been reported is insufficient to alter the likelihood or latency of cue-elicited reward seeking.

While the largest increases in VP→VTA calcium activity consistently occurred following port-entry, smaller peaks were observed in some subjects ∼1s after DS onset. What might activity in the time epoch between cue onset and port-entry represent, and why does this vary between subjects? Calcium dynamics in this time epoch may be related to individual differences in incentive value attribution to the DS between-subjects. Indeed, individual differences in cue-evoked VP neuronal activity have been observed between individuals exhibiting “sign-tracking” and “goal-tracking” behavior in response to Pavlovian cues (Ahrens et al., 2016), and changes in incentive value of cues has also been associated with changes in VTA dopaminergic signaling (Berridge and Aldridge, 2014; Koob and Volkow, 2016; Volkow et al., 2017). These differences are well-documented in Pavlovian paradigms with visual cues, but their expression varies with cue properties (Meyer et al., 2012, 2014) so careful experimental design is required for their detection. It is unclear whether individual differences in incentive value attribution to cues manifest similarly in instrumental paradigms such as the DS task. Future work should investigate the generalizability of our VP→VTA findings to Pavlovian cues, and even visual instrumental cues.

### Role of VP→VTA neurons in motivated behavior

Our results suggest that VP→VTA neurons are reinforcing, but do not contribute to response invigoration by cues and reward. We report optogenetic stimulation of VP→VTA neurons supports ICSS in rats, consistent with prior work in mice showing activation of GABAergic VP→VTA neurons is reinforcing (Faget et al., 2018). There were noteworthy differences in the magnitude of ICSS effects between-subjects within the VP→VTA group. Though we did not observe a clear pattern of subregional (rostral versus caudal or medial versus lateral) implant placements associated with stronger effects here, subregional differences in VP function have been reported (Smith et al., 2009; Mahler et al., 2014; Root et al., 2015). Additionally, some variability in our photometry and behavioral findings may be due to differential recruitment of GABAergic versus glutamatergic neurons. While the majority of VP neurons are GABAergic (Smith et al., 2009), the viral approach used in this study did not target a specific neuronal subpopulation and prior work demonstrates that GABAergic and glutamatergic VP neurons have functionally opposing activity patterns *in-vivo*, generally associated with approach and avoidance respectively (Faget et al., 2018; Stephenson-Jones et al., 2020).

Our observation of robust VP→VTA calcium activation during sucrose consumption, coupled with our findings that optogenetic stimulation of VP→VTA neurons paired with sucrose consumption biases choice, suggests that VP→VTA neurons may encode some aspect of the value of reward consumed and reinforce seeking behavior accordingly. From a circuit perspective, reinforcement may occur through disinhibition of dopaminergic VTA neurons from VTA GABAergic interneurons by reward-evoked GABAergic VP→VTA activity (Hjelmstad et al., 2013; Faget et al., 2018). GABAergic VTA neurons are not homogeneous and GABAergic VTA projections have themselves been associated with motivated behavior (Morales and Margolis, 2017). Recent work shows that GABAergic neurons projecting from the VTA to the VP encode information about reward value and that optogenetic stimulation of these projections enhances motivation to seek reward without providing robust reinforcement (Zhou et al., 2022). Thus, VP and VTA seem to provide reciprocal feedback about the value of rewards consumed to support learning and efficient reward capture.

Interestingly, we found that optogenetic stimulation of VP→VTA neurons paired with sucrose consumption biases choice without impacting the hedonic value of sucrose, as measured by licking behavior. This finding contrasts with other work demonstrating that optogenetic stimulation of ventral arkypallidal projections to the nucleus accumbens shell paired with sucrose consumption enhances the hedonic value of sucrose (Vachez et al., 2021), reaffirming the diverse functional roles of VP output pathways. Our data suggest that VP→VTA neuron activation may bias seeking behavior independently of experienced outcome value. This potential disconnection between seeking behavior and hedonic value is noteworthy, since compulsive seeking despite negative affective outcomes observed in addiction is associated with changes in VTA dopaminergic signaling (Robinson and Berridge, 2000; Koob and Volkow, 2016; Volkow et al., 2017). Since animals only received liquid sucrose reward in our experiments, the relationship between VP→VTA activity and outcome representation more generally, including other consummatory rewards, aversive outcomes, and non-consummatory outcomes, requires further study.

### Reconciling these data with prior reports of VP→VTA circuit roles in cued reinstatement

Past work investigating the role of VP→VTA neurons in cue-elicited reward seeking (Mahler and Aston-Jones, 2012; Mahler et al., 2014) has largely been limited to cue-induced reinstatement paradigms which test the ability of action-contingent presentations of a Pavlovian drug-conditioned stimulus to reinstate extinguished drug-seeking behavior. In other words, animals perform operant seeking behavior to “earn” drug cue presentations in the absence of drug. Thus, reinstatement of extinguished behavior observed in these paradigms may not reflect drug-seeking or craving *per se* but may instead be due to conditioned reinforcement by drug cues (Troisi, 2013; Namba et al., 2018; Lay and Khoo, 2021). Though we did not observe a consistent DS response here, contingent delivery of Pavlovian drug cues in past studies may have elicited sufficient VP→VTA activation to reinforce seeking behavior. Other work has investigated VP→VTA contributions to non-contingent drug “cues” in a context renewal paradigm (Prasad et al., 2020), but this work used a serial chemogenetic disconnection approach which may have also disrupted the reciprocal projections from VTA to the VP, or bisynaptic connectivity via other structures (Root et al., 2015; Haber, 2017; Morales and Margolis, 2017). Thus, the degree to which VP→VTA neurons contribute to context-induced relapse remains unclear. Since the relapse models used in prior work test reinstatement of seeking behavior following extinction, it’s also possible that durable changes in VP→VTA activity occur during extinction learning or in the absence of drugs which make animals susceptible to relapse. This highlights a need for more basic understanding of how VP→VTA circuit function changes with experience and reflects internal states.

### Effects of VP→mdThal optogenetic stimulation

Relative to many other VP output pathways, the role of VP→mdThal in motivated behavior is not well understood. This pathway is broadly implicated in consolidation of action-outcome contingencies (Mitchell and Chakraborty, 2013; Soares-Cunha and Heinsbroek, 2023). Prior work has reported dissociable roles of VP→mdThal and VP→VTA neurons in Pavlovian-to-instrumental transfer (PIT) (Leung and Balleine, 2015), a phenomenon thought to reflect the incentive value of cues in which noncontingent Pavlovian cue presentations increase instrumental reward seeking behavior despite the absence of the primary reinforcer. Similarly, we report dissociable effects of optogenetic manipulations of these pathways. Notably, we found that optogenetic stimulation of VP→mdThal neurons did not support ICSS. This suggests that, while this pathway may be important for reward learning, it is not directly reinforcing. In addition, we found that optogenetic stimulation of VP→mdThal neurons paired with sucrose consumption did not impact choice behavior in the lever choice task, in contrast to our VP→VTA findings. Finally, we did not find any impact of cue-paired optogenetic stimulation of VP→mdThal neurons on probability or vigor of seeking behavior in rats trained in the DS task. Since these manipulations occurred in well-practiced rats, this finding is consistent with prior reports that mdThal is important for consolidation, but not the expression of learned instrumental reward seeking (Mitchell and Chakraborty, 2013). Altogether our results suggest that cue-paired VP→mdThal neuronal activity is not sufficient to promote generalized seeking in response to cues, and is insufficient to alter outcome representations or bias instrumental action selection. More work is required to investigate mechanisms by which VP→mdThal contributes to outcome-guided behavior.

### Conclusions and Future Directions

Taken together, results from this study suggest that VP→VTA neurons are active during the consumption of natural rewards and that this activity reinforces seeking behavior. Critically, we found that VP→VTA population-level calcium signals do not encode reward-seeking vigor in a discriminative stimulus task, and that cue-paired VP→VTA optogenetic stimulation in this task does not impact the vigor of reward seeking. Thus, a key question remains: If not VP→VTA neurons, what VP single units encode vigor of instrumental responding to cues (Richard et al., 2016, 2018; Lederman et al., 2021)? Photometry recordings collected in our lab revealed GABAergic VP neuron calcium activity encodes response vigor in the DS task (Scott et al., 2022). There may be no single VP “output pathway” responsible for controlling vigor, but coordinated recruitment of GABAergic VP neurons is critical for controlling animals’ motivational state (Faget et al., 2018; Stephenson-Jones et al., 2020). Future work interrogating the contributions of more specific VP→VTA cell types (Hjelmstad et al., 2013; Knowland et al., 2017; Faget et al., 2018; Prasad et al., 2020) or individual neuronal responses may reveal more nuanced roles of these neurons in cue-elicited reward seeking.

